# Nitrate fertilization may delay autumn leaf senescence, while amino acid treatments do not

**DOI:** 10.1101/2021.11.09.467959

**Authors:** Nazeer Fataftah, Erik Edlund, Jenna Lihavainen, Pushan Bag, Lars Björkén, Torgny Näsholm, Stefan Jansson

**Author notes:** Corresponding author: Stefan Jansson. these authors contributed equally to this work (Nazeer Fataftah, Erik Edlund). **Author for contact:** Stefan Jansson, Umeå Plant Science Centre, Department of Plant Physiology, Umeå University, Umeå, Sweden. The author responsible for distribution of materials integral to the findings presented in this article in accordance with the policy described in the Instructions for Authors (https://academic.oup.com/plphys/pages/General-Instructions) is Stefan Jansson. **Author contribution** NF, EE, and SJ conceived the study, SJ and TN supervised the experiments; NF, EE and LB designed and performed the fertilization experiment and analyzed the data. NF and JL performed the metabolomics analyses. PB was responsible for the chl data fitting and preparing figures. NF, EE, JL, and SJ wrote the article and all authors discussed and commented on it. SJ is the author responsible for contact and ensures communication.

## Abstract

Fertilization with nitrogen (N)-rich compounds leads to increased growth, but may compromise phenology and winter survival of trees in boreal regions. During autumn, N is remobilized from senescing leaves and stored in other parts of the tree to be used in the next growing season. However, the mechanism behind the N fertilization effect on winter survival is not well understood and it is unclear how N levels or forms modulate autumn senescence. We performed fertilization experiments and showed that treating *Populus* saplings with high or low levels of inorganic nitrogen resulted in a delay in senescence. In addition, by using precise delivery of solutes into the xylem stream of *Populus* trees in their natural environment, we found that delay of autumn senescence was dependent on the form of N administered: inorganic N (NO_3_^−1^) delayed senescence but amino acids (Arg, Glu, Gln, and Leu) did not. Metabolite profiling of leaves showed that the levels of tricarboxylic acids (TCA), arginine catabolites (ammonium, ornithine), glycine, glycine-serine ratio and overall carbon-to-nitrogen (C/N) ratio were affected differently by the way of applying NO_3_^−1^ and Arg treatments. In addition, the onset of senescence did not coincide with soluble sugar accumulation in any of the treatments. Taken together, metabolomic rearrangement under different N forms or experimental setups could modulate senescence process, but not initiation and progression in *Populus*. We propose that the different regulation of C and N status through direct molecular signaling of NO_3_^−1^ could account for the contrasting effects of NO_3_^−1^ and Arg on senescence.

**One sentence summary:** Nitrate, administered by precision fertilization through injection into the trunk, may delay autumn senescence and change metabolism in *Populus* leaves, while the same amount of amino acids does not have the same effect.

## Introduction

Plants have evolved to sense variation in abiotic factors and adjust their development based on this information. Such acclimation is essential for fitness under variable conditions, but variation in the environment also leads to adaptation, which is the evolutionary process that modifies genetic makeup to make the plant better able to live in its habitat. For plants in general, and trees in temperate regions in particular, phenological traits are extremely important; inability to respond to seasonal cues inducing the annual cycle of growth and rest would be detrimental (Dox et al, 2020). Leaf senescence can be initiated by developmental age or unsuitable environmental conditions such as nutrient deficiency, but deciduous trees in temperate regions have typically developed the ability to initiate senescence at a given time in the autumn, presumably driven by light cues (Michelson et al. 2018, Fataftah et al., 2021). This autumnal senescence is a tightly controlled developmental process during which many cell components are degraded and, for example, nitrogen (N) is remobilized from the senescing leaves and stored over the winter in other tree parts such as in bark and roots, allowing it to be used during the next growing season (Keskitalo et al., 2005, Millard and Grelet, 2010). Most forest soils are strongly depleted in these nutrients, trees that have adapted to correctly time their autumnal senescence, therefore, have a strong competitive advantage. Trees that do not complete the senescence process before the leaf cells are killed by harsh environmental conditions such as frost resorb fewer nutrients and can produce less foliage the following season and be less competent to harvest sunlight. On the other hand, trees that enter senescence too early loose valuable photosynthates, which in a climate with a short vegetative season can be detrimental for long-term survival. Thus, the timing of autumn senescence in temperate regions represents a trade-off between N retention and carbon (C) acquisition. In other climates, avoiding resources being lost as a result of factors such as wind, drought, or pathogens could be more important for the regulation of leaf senescence.

It is a “common gardening practice” not to apply N-rich fertilizer to trees in late season as this may compromise winter survival; however, the physiological basis for this is not well understood. N can be present in the soil in different forms; commercial N fertilization of agricultural plants and scientific studies on plant fertilization typically use inorganic N in the form of ammonium and nitrate, or in an organic form such as urea. However, there is a growing body of evidence that in particular in the forest ecosystem, but probably also in other ecosystems where commercial fertilizers are not used, N is mainly available as forms of organic N, predominantly amino acids (Näsholm et al. 1998, Ganeteg et al. 2017, Schulten et al., 1998, Yu et al., 2002, Andersson et al., 2004), and that organic N may contribute more to plant N supply than often assumed (Brackin et al. 2015, Czaban et al. 2016, Farzadfar et al, 2021).

We postulate that *Populus* leaves have to acquire “competence to senesce” before they can respond to autumnal cues that induce senescence (Fracheboud et al. 2009). As autumn senescence is a trade-off between N and carbon metabolism, we reason that N status could affect autumn senescence and have designed a series of experiments addressing the hypotheses that 1) N levels influence autumn senescence and 2) inorganic and organic forms of N differ in their capacity to influence the autumn phenology in *Populus.*

## Results

### Fertilization may override the control of autumn senescence in *Populus*

To study the effect of nutrient fertilization on the progression of autumn senescence, we established a small common garden experiment with four natural aspen genotypes from the SwAsp collection (Luquez et al. 2007), originating from the same latitude as the common garden. The trees were fertilized with a commercial fertilizer, but once all trees had set bud in July some were continuously treated (high fertilizer; HF) while no additional fertilizer was given to the other half (low fertilizer; LF). One month after the separation of the fertilization regimes, leaves of HF trees had more than twice the amount of chlorophyll compared to LF treated trees (Figure 1Ai). Senescence, defined as an accelerated chlorophyll degradation, started between August 25^th^ and 27^th^ for all genotypes in the LF treatment (Figure 1Ai). This senescence timing fitted with the data from a nearby common garden (which has not received fertilization for several years) – where these genotypes initiate senescence between August 27^th^ and September 12^th^ (Fracheboud et al. 2009). In addition, anthocyanin was highly accumulated in the LF treatment (Figure 1Aii; B). In the HF treatment, trees showed accelerated chlorophyll degradation starting about a month later than in the LF treatment (Figure 1Ai). Once initiated, the chlorophyll degradation process was rapid and accelerated in the first week of October, presumably due to frost damage. Leaves of the HF trees did not accumulate anthocyanins at all, unlike the leaves in the LF treatment; instead anthocyanin levels decreased in advanced stages of senescence (Figure 1Aii). HF resulted, therefore, in very different leaf pigmentation: leaves sampled on the same day from different genotype/treatment combinations are shown in Figure 1B.

**Figure 1.**
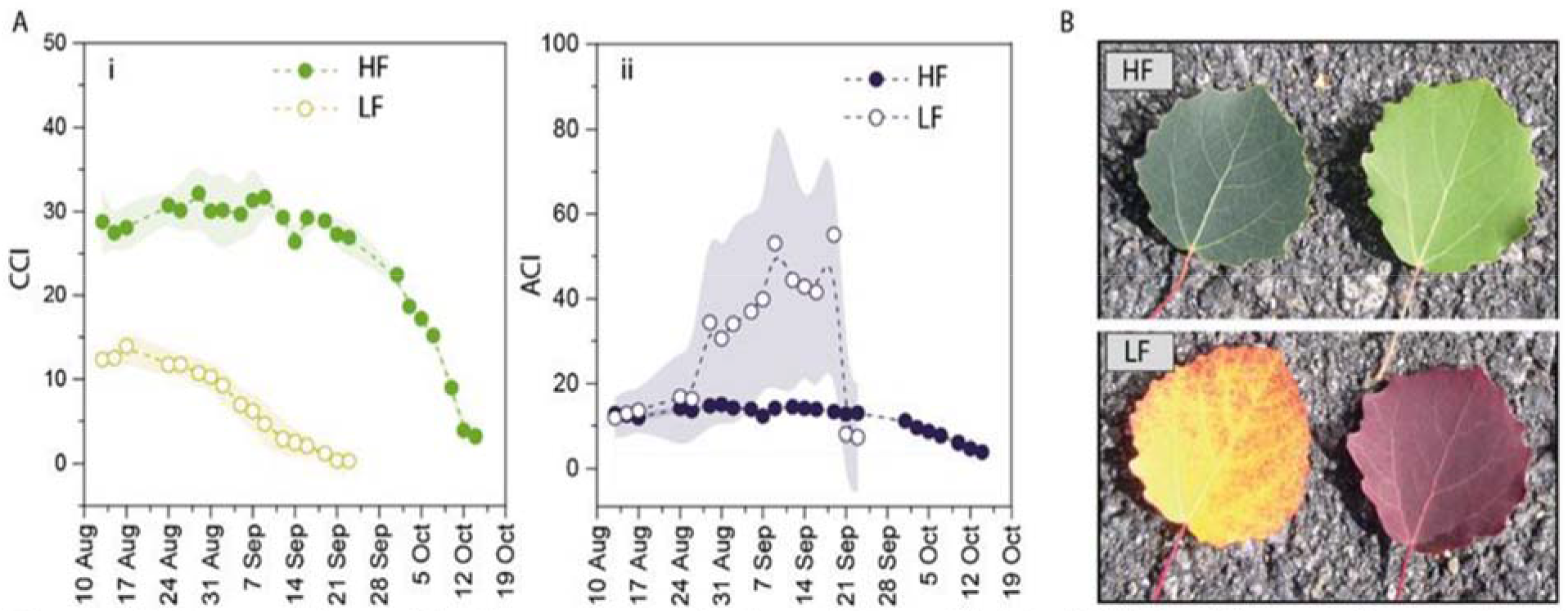
Progression of leaf senescence low (LF) and high (HF) fertilizer regimes. Ai and Aii, chlorophyll and anthocyanin content indices of the mean of four replicated genotypes, respectively, under low fertilizer (LF) and high fertilizer (HF) regimes grown in a common garden during autumn 2008. B, leaf coloration of the aspen genotypes in LF and HF treatments. Photo was taken on September 2^nd^ 2008.

We next investigated the effect of N availability on autumn senescence by only changing the N fertilization level while keeping the levels of the other nutrients constant. Hybrid *Populus* (T89) saplings were treated with a nutrient solution containing all required nutrients including 9 mM NH_4_NO_3_ (HN) or no additional NH_4_NO_3_ (LN) in the greenhouse. On August 22^nd^, the saplings were transferred to outdoor conditions, the photoperiod at this date is 15.5 hours corresponding to the critical daylength for bud set for this particular genotype. On September 5^th^, the trees of both treatments were of similar height, indicating that the treatment was not harsh enough to have dramatic effects on growth, and lower leaves of HN and LN, developed early in the HN/LN treatments, had similar chlorophyll levels (Figure 2A-Ci). However, the HN leaves that developed during the treatments contained ca. 50 % more chlorophyll than LN leaves (Figure 2Cii). Chlorophyll levels started to diverge between the treatments; in particular, in the lower leaves of LN plants, chlorophyll content had already started to decrease at the end of August (Figure 2Ci). However, chlorophyll levels in the lower and upper leaves of HN saplings started to decrease around the same time as each other, on September 25^th^ (Figure 2C). Although the upper leaves of LN saplings had approx. 50% lower chlorophyll levels compared to HN leaves, they also started to senescence at the end of September (Figure 2Cii). This suggests that the developmental stage of the leaves and relative changes in N levels during tree development could account for early senescence onset in lower leaves in the LN treatment. Taken together, these two experiments show that fertilization could override the normal control of autumnal senescence in *Populus* and delay it.

**Figure 2.**
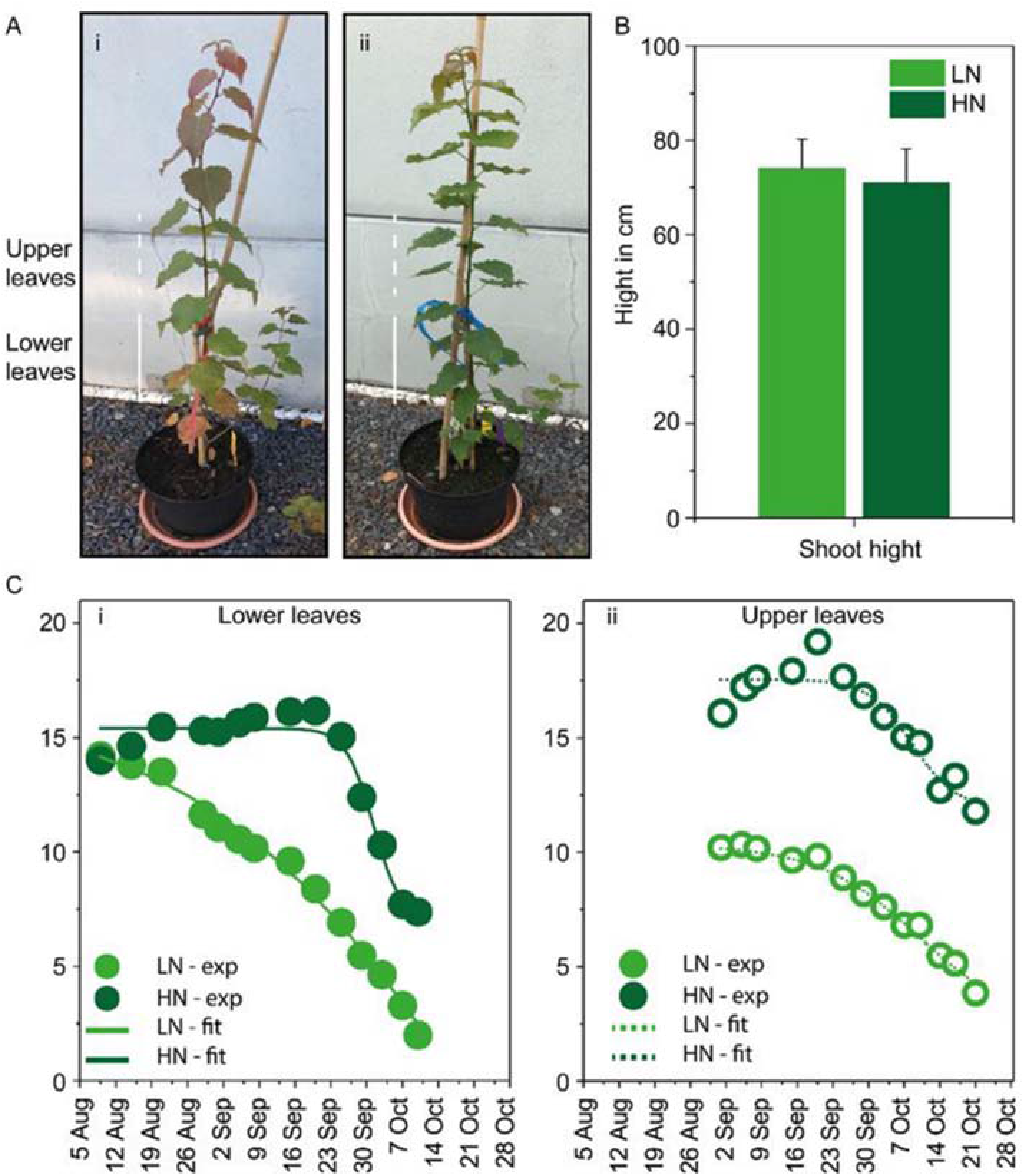
Progression of leaf senescence and growth under high or low N regimes. A, phenotype of hybrid aspen saplings under low (LN) (A-i) and high (HN) (A-ii) nitrogen regimes; continuous and dotted lines indicate the regions for lower and upper leaves, respectively. B, shoot height of saplings was measured on 5^th^ of September 2018 in LN and HN treatments; bars represent mean ± SD, n=6. C, chlorophyll content index (CCI) of leaves from the lower part of the shoot (C-i) and from the upper part of the shoot (C-ii); dots represent mean CCI values of six trees and the lines represent the fitted curves.

### Nitrate, but not amino acid, injection into the xylem delayed autumn senescence

Soil fertilization with NH_4_NO_3_ is able to influence the onset of autumn senescence in small trees and saplings. How relevant is this for trees in their natural environment – a forest soil – where N is mainly available in its organic form mainly as amino acids (Yu et al., 2002, Andersson et al., 2004, Inselsbacher and Näsholm 2012)? Much *Populus* propagation in the natural environment is through clonal replication, with several suckers emerging from the same root, establishing mature trees sharing the same root system. This opens up the possibility to expose genetically identical tree trunks sharing a root system to different treatments within a naturally established tree stand. We used a modification of the liquid injection system (Swanston and Myrold, 1998), in which an infusion bag is connected to the trunk through a syringe in the injection hole, allowing for the addition of solutes passively into the xylem stream of individual tree trunks (Figure 3A). Using this system, we added controlled amounts of either inorganic N as nitrate (NO_3_^−1^) or organic N as arginine (Arg). We gave four different doses of N on eight occasions over the season, in total 4, 8, 20, or 60 g of N per trunk were injected (hereafter named N4, N8, N20, and N60 for NO_3_^−1^, and A4, A8, A20 and A60 for Arg treatments). All trees appeared healthy at the end of the treatments. Data from the three stands were analyzed both separately and combined; here we show the data from the three stands combined (Figure 4), the chlorophyll data for individual stands can be found in Supplementary Fig S1.

**Figure 3.**
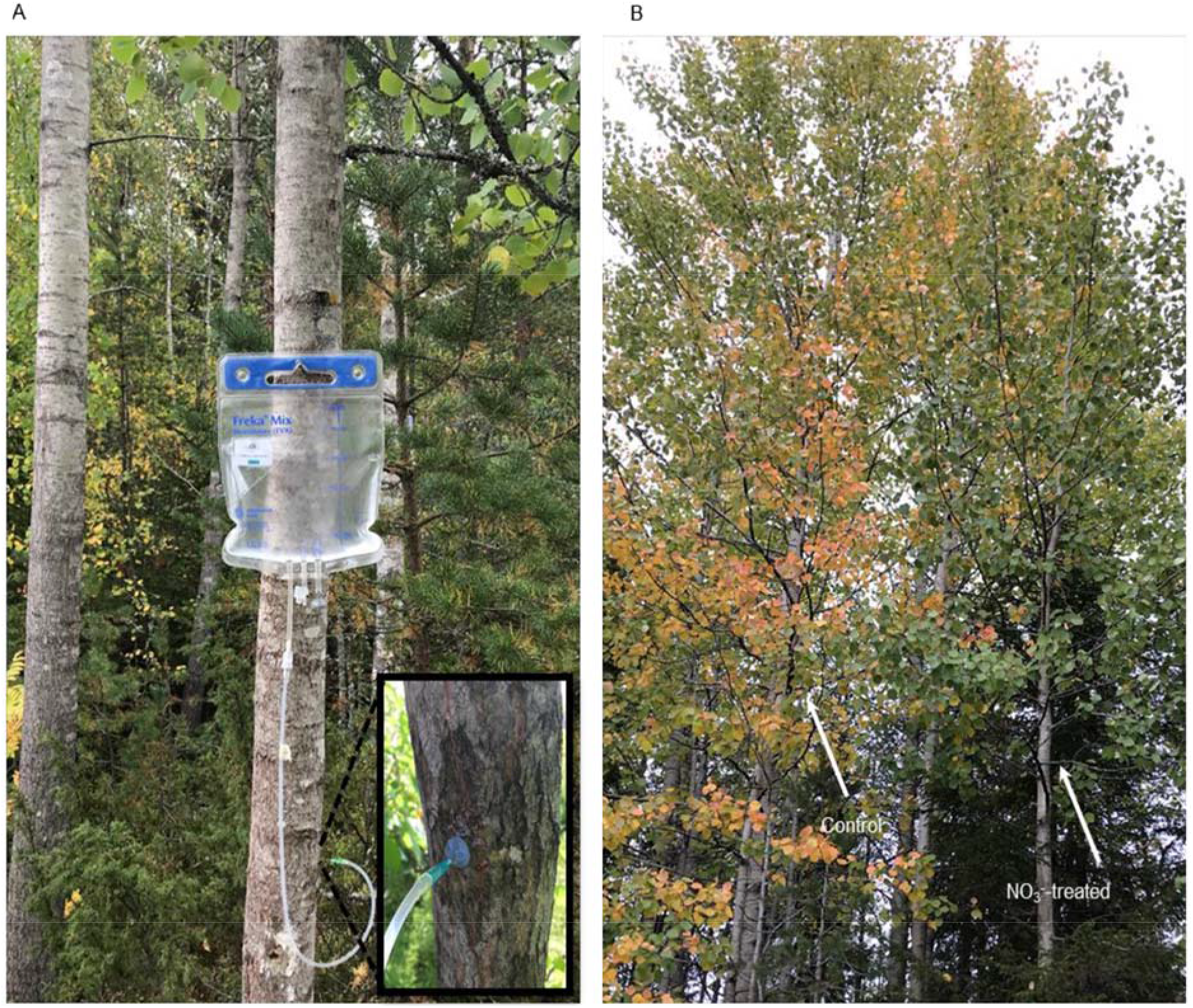
Xylem fertilization treatment setup and its effect on autumn phenology in aspen. A, experimental setup of injection application to aspen trunks with medical IV (intravenous) bags connected to the xylem through a syringe. B, the effect of NO_3_^−1^ application on aspen autumnal leaf senescence progression; control tree on the left was injected with water and the tree on the right was injected with NO_3_^−1^ in late autumn.

**Figure 4.**
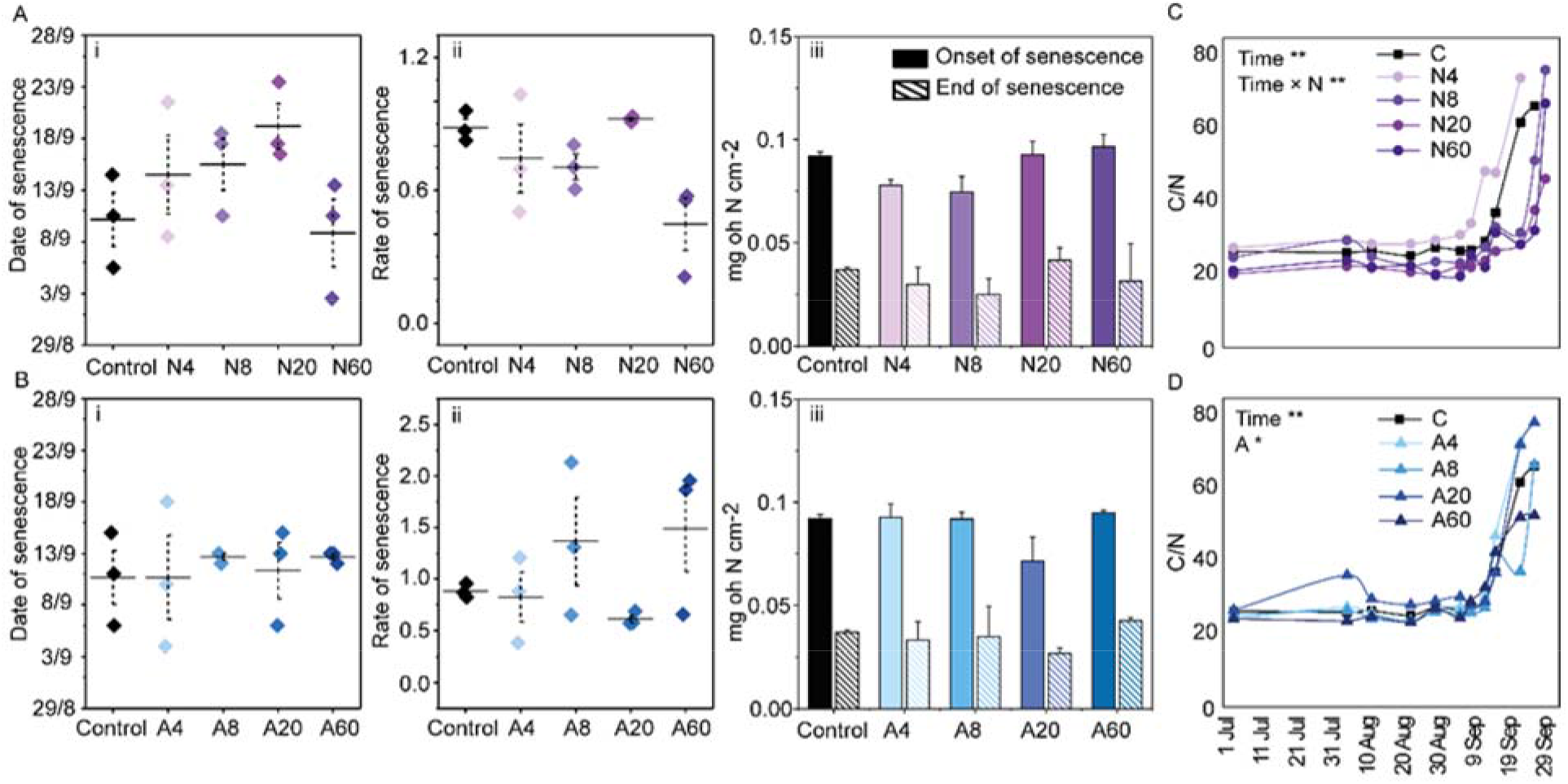
Effects of xylem fertilization with nitrate and arginine on autumn leaf senescence in aspen. A and B, progression of autumnal leaf senescence in 2011 in aspen trunks injected with water (Control), and various amounts of NO_3_^−1^ or arginine, respectively. Date of senescence onset (i), rate of senescence (ii), and the leaf nitrogen levels of N at the start and end of senescence (iii); Dots represent original data points from three replicate trees, solid line represents the mean and dashed line represents ±SD; n=3. Bars represent mean of three trees ± SD (iii).

Neither the NO_3_^−1^ nor Arg treatments resulted in a consistent change in pre-senescent N or chlorophyll levels of the leaves (Figure 4Aiii; Biii, Supplementary Figure S2). However, nitrate fertilization seemed to result in delayed senescence, apparently in a dose-dependent manner, although N60 trees senesced at the same time as control trees (Figure 3B; Figure 4Ai). Fertilization with Arg did not, on average, affect senescence progression (Figure 4B), although in one stand Arg slightly delayed it (Supplementary Figure S1). Moreover, xylem fertilization treatment with NO_3_^−1^ delayed the increase in C/N ratio (except in N4, Figure 4C), which typically occurs as a result of activated N remobilization in senescing leaves. Arg injection, on the other hand, did not delay the increase in C/N ratio, on the contrary in A20 it appeared earlier (Figure 4D). However, neither of the treatments affected the remaining N content at the end of senescence (Figure 4Aiii; Biii).

In another year – and in other stands – we repeated the experiment with NO_3_^−1^ and other organic N forms using a similar setup. The N solutions included inorganic N (NO_3_^−1^), or organic N as amino acids (Glu, Gln, or Leu). As the three stands had similar pre-senescence chlorophyll levels, we combined the data from them (Figure 5). Consistent with the earlier experiment, the addition of a small amount of N (5 g) as NO_3_^−1^ delayed the onset of senescence, whereas organic nitrogen species (Glu, Gln, or Leu) had no effect (Figure 5). Taken together, these results showed that addition of nitrate ions to the xylem sap delayed senescence in *Populus*, while addition of amino acids did not.

**Figure 5.**
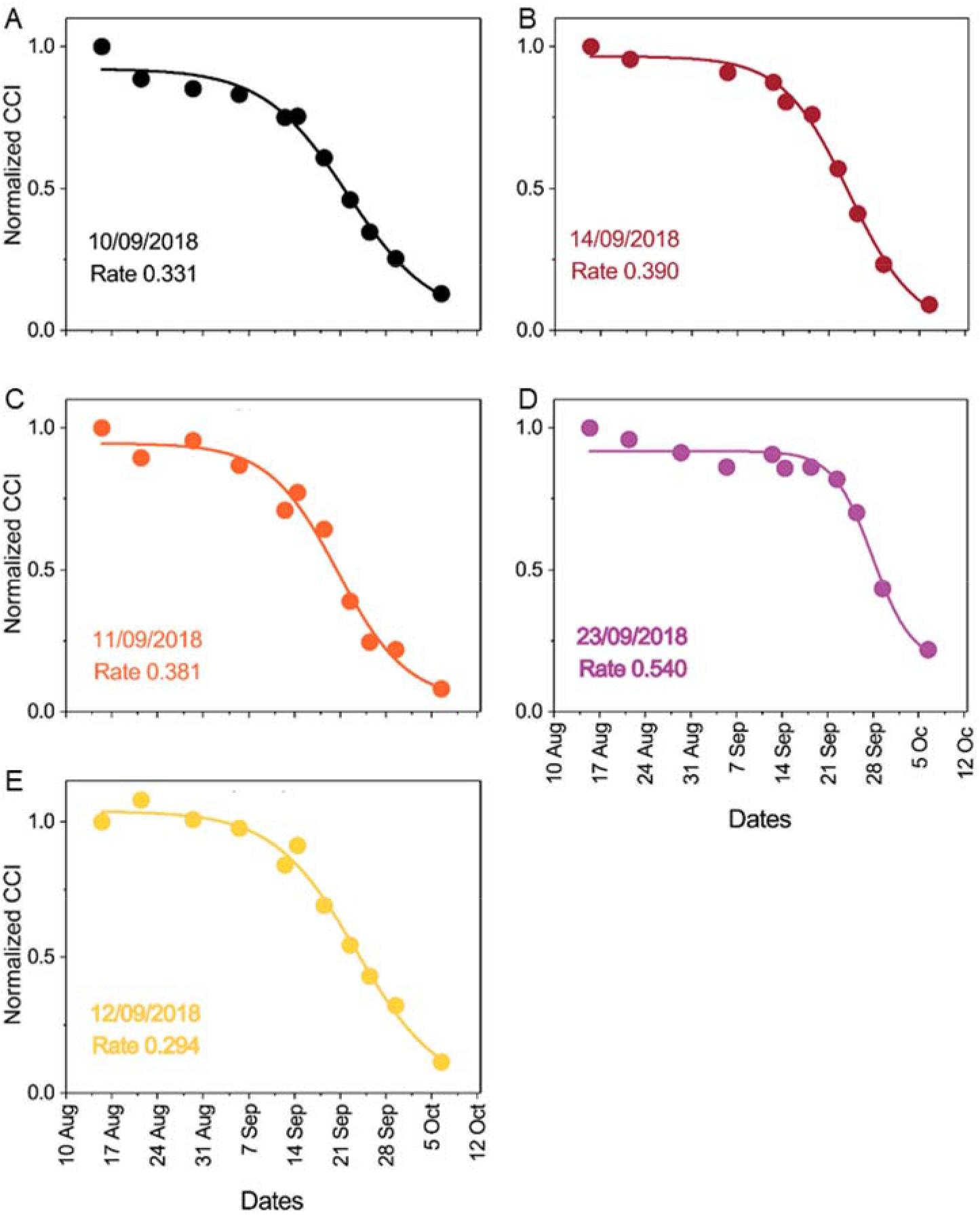
The effect of inorganic nitrogen and amino acid injections on the onset and rate of autumnal senescence in aspen in 2018. Senescence onset was defined based on the start of rapid chlorophyll depletion. CCI values were expressed relative to the starting values on August 12^th^. Control trunks were injected with water (A) and glutamine (B), glutamate (C), KNO_3_ (D) or leucine (E). Dots represent the mean value of three trees (n=3), and the lines represent the fitted curves. Date of senescence onset and senescence rate are represented in the plots.

### Soil nitrogen fertilization resulted in a metabolic shift in the leaves of hybrid *Populus*

These experiments demonstrate that nitrate ions may have a signaling function that could delay the onset of senescence in *Populus*. We performed metabolite analyses on the lower leaves from trees in the soil fertilization experiment with the HN treatment or without additional NH4NO3 (LN, Figure 2) in autumn 2018. Principal Component Analysis (PCA) revealed that the metabolite profiles in leaves in low N (LN) and high N (HN) treatments already differed at the first time point (August 16^th^) based on the first principal component (PC1 20%, Figure 6A). The second component explained the variation in the data across the four time points (PC2 17%, Figure 6A). Out of the 86 annotated metabolites, the levels of 46 individual metabolites as well as Gln/Glu (Glutamine-to-Glutamic acid) and Gly/Ser (Glycine-to-Serine) ratios, indicators of ammonium assimilation and photorespiration, respectively, were significantly affected by the N treatments (Figure 6B, Supplementary Table S1). The most important metabolites to account for the separation of the N treatments were identified based on a volcano plot (Figure 6C). Metabolites with higher concentrations in HN than in LN included amino acids (Ser, Gln, Asp, Ala, and Lys) and the metabolites at lower levels included tricarboxylic acids (TCA), malic acid, isocitric acid and citric acid (Figure 6C). The higher Gln/Glu ratio suggests that ammonium assimilation was enhanced and the lower Gly/Ser ratio that photorespiration was repressed in HN plants compared to LN plants (Figure 6B). The accumulation of soluble carbohydrates has generally been associated with the onset of senescence, especially under N deficiency. However, unexpectedly, the levels of hexoses – glucose, fructose and galactose – were higher in HN than in LN irrespective of time point (Figure 6C, Figure 7). Therefore, we conclude that the differences in senescence behavior between LN- and HN- treated trees were not associated with soluble sugar accumulation, rather it appeared that they could be related to altered direct NO_3_^−1^ signaling and/or TCA cycle function.

**Figure 6.**
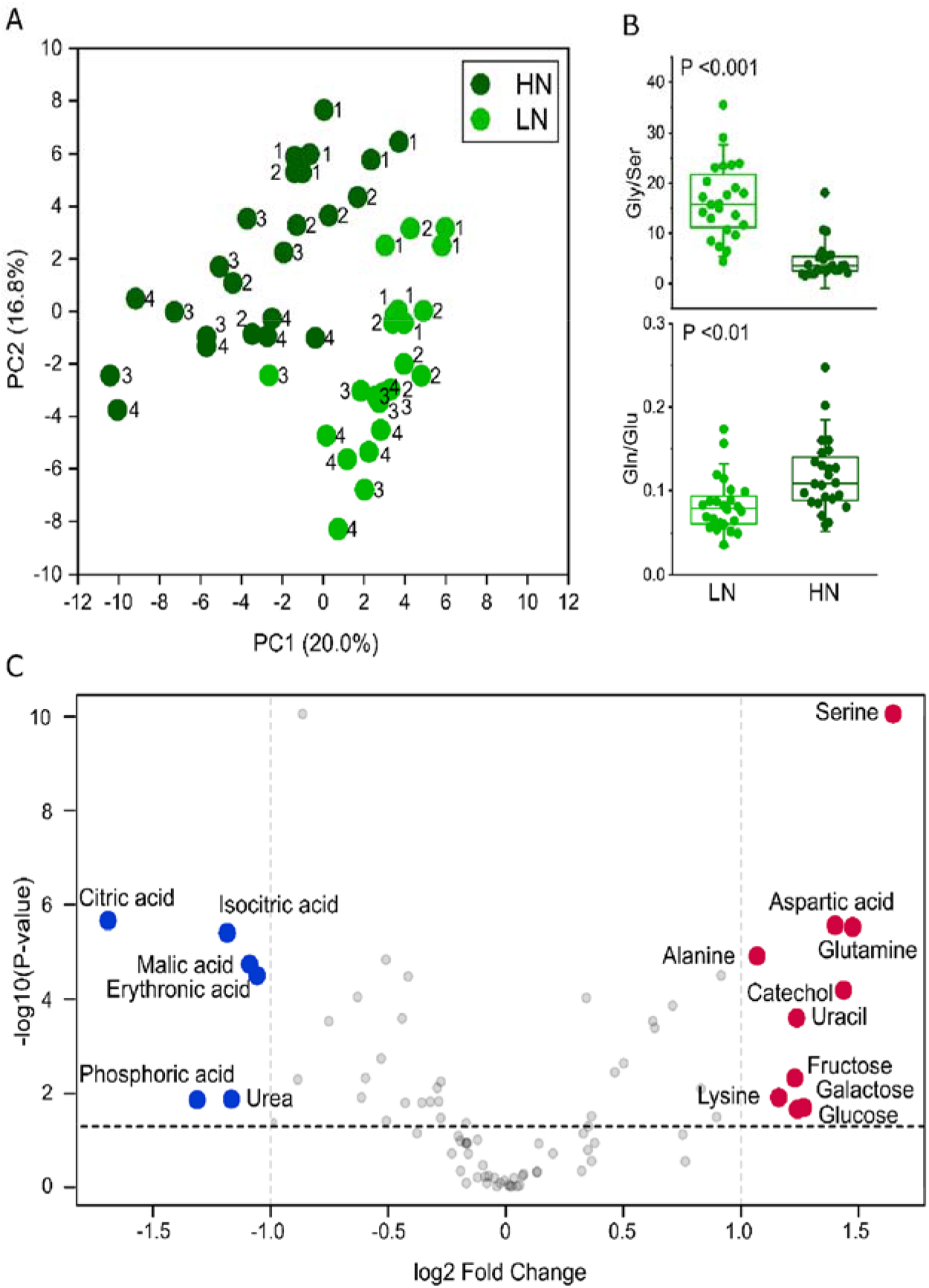
Leaf metabolite profiles under different nitrogen regimes. A, principal component analysis (PCA) revealed variation in the metabolite profiles (86 metabolites) between the LN and HN treatments based on the first principal component (PC) irrespective of time point (*n*=6 for each time point per treatment); Time points were separated based on the second principal component (PC2). B, glycine-to-serine (Gly/Ser) and glutamine-to-glutamic acid (Gln/Glu) ratios as indicators of photorespiration and ammonium assimilation, respectively, were affected by the nitrogen treatments; the effect of nitrogen treatment was tested with a t-test and the box plot shows original data points, the box represents 25% and 75% with the mean, and whiskers ± SD, *n*=24 for each treatment. C, volcano plot showing the effect of nitrogen treatment on metabolite levels; Metabolites with higher and lower abundance in HN than in LN are in red and blue, respectively (log_2_ fold change cut off >1.0, FDR adjusted *P*-value <0.05, t-test). Details of statistical results are in Supplementary Table S1.

**Figure 7.**
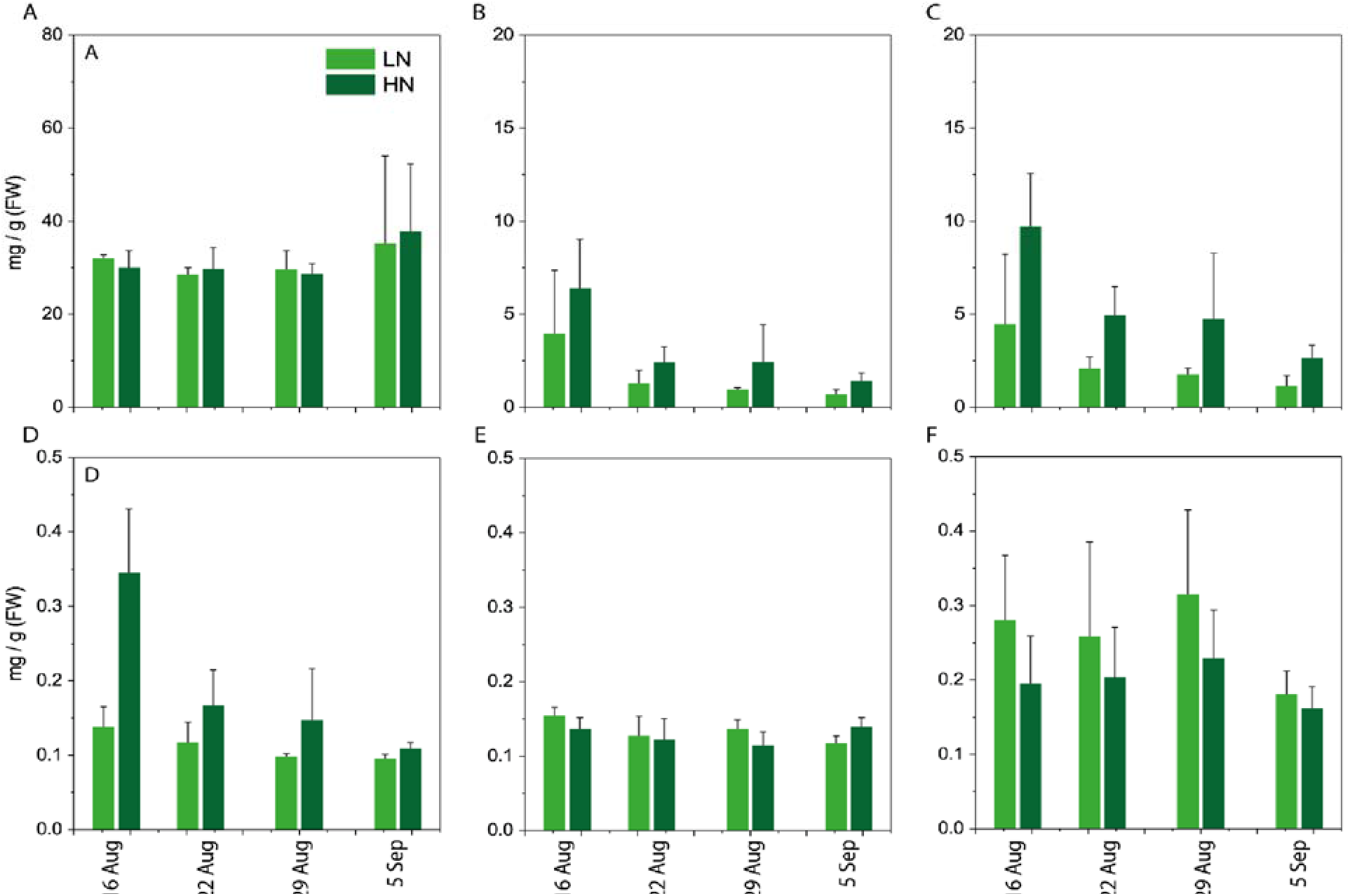
Carbohydrate levels under LN and HN regimes during autumn. A-F, the levels of sucrose (A), glucose (B), fructose (C), galactose (D), trehalose (E) and maltose (F) were quantified (mg/g fresh weight) at four time points during autumn 2018 in trees growing under LN and HN regimes. Bars indicate means ± SD (n= 3).

### The metabolic shift was more pronounced in nitrate than in arginine treatments

Finally, we investigated how various injected concentrations of NO_3_^−1^ or Arg affected the metabolite profiles of the leaves (sampled during the experiment presented in Figure 4). As the foliar metabolite profiles were initially different for the individual trunks, the levels were normalized with respect to samples collected at the first time point before the xylem fertilization started. The first principal component (PC1) explained 18.0% of the variation in the data and was associated with time (Figure 8A). The third (PC3 9.0%) and the fourth (PC4 7.4%) principal components separated the injection treatments (Figure 8B). Overall, NO_3_^−1^ treatment had a stronger effect on the metabolite profile than the Arg treatment; all NO_3_^−1^ treatments and the two highest Arg treatments (A20, A60) were separated from the control and the other Arg treatments (Figure 8B). Out of the 86 annotated metabolites, the levels of 36 were significantly affected by time irrespective of the treatment, 26 by NO_3_^−1^, and 23 by Arg treatment (Figure 8C, Supplementary Table S2 and S3).

**Figure 8.**
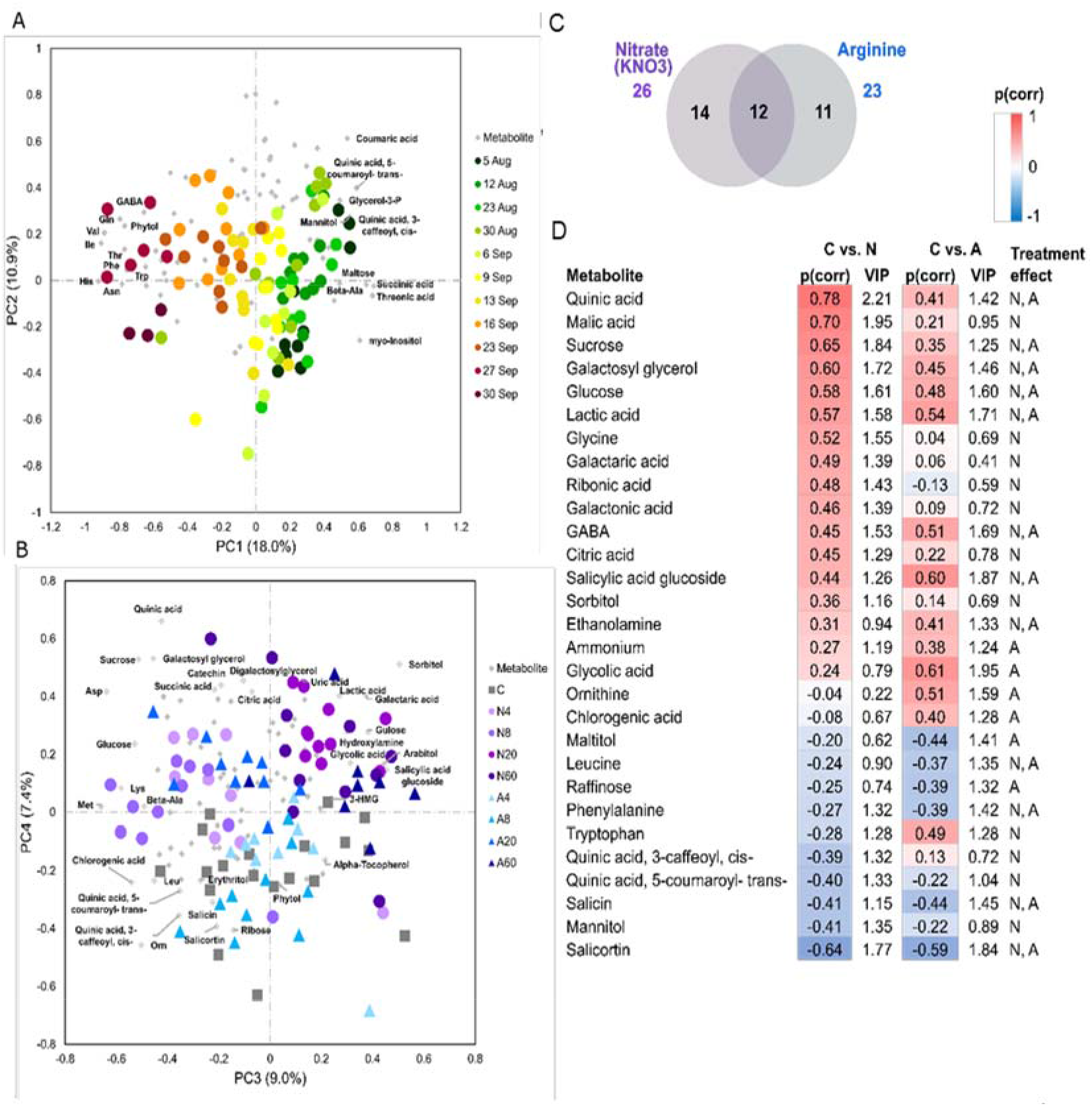
Leaf metabolite profiles in aspen as affected by xylem fertilization with NO_3_^−1^ and arginine in autumn. A, biplots of principal component analysis (PCA) of leaf metabolite data (86 metabolites) in xylem fertilization experiments in autumn 2011 revealed that the first principal component (PC1) was associated with variation in the metabolite profile based on time. B, the third (PC3) and fourth (PC4) principal components separated the treatments; Metabolites accounting for most of the separation of time points and injection treatments are labelled. Metabolite data were normalized based on the pre-fertilization level on July 4^th^, *n*=3-10 at each time point (A), *n*=7-18 for each treatment (B). C, leaf carbon-to-nitrogen ratio (C/N) as affected by xylem fertilization treatments during autumn 2011 in aspen leaves. The effects of time, treatments (nitrate [N] or arginine [A] fertilization) and their interaction with time on C/N ratio were tested with two-way ANOVA, *p*-value *<0.05, **<0.01. D, Venn diagram depicts the overlap and specific metabolites affected by nitrate and arginine treatments. E, discriminant analysis (OPLS-DA) was performed to identify the metabolites with high p(corr) >|0.3| and VIP-scores >1.0, accounting the most for the separation of nitrate (C vs. N) or arginine (C vs. A) treatments from the control. Treatment effects on metabolite levels were tested with ANOVA, FDR adjusted *P*-value <0.05 considered significant. Details of statistical results are in Supplementary Tables S2 and S3.

Metabolite responses to NO_3_^−1^ and arginine treatments were compared, and discriminant analyses (OPLS-DA) were performed in order to identify the metabolites associated with the greatest treatment effects (Figure 8D). The levels of 12 metabolites were similarly affected by both treatments (Figure 8D). The levels of sucrose, glucose, lactic acid, GABA, and salicylic acid glucoside were higher, whereas the levels of salicin, salicortin, leucine, and phenylalanine were lower in NO_3_^−1^ and Arg treatments than in the control (Figure 8D). Metabolite ratios related to pyruvate metabolism responded similarly to both forms of N; lactate-to-pyruvate (Lac/Pyr) and malate-to-pyruvate (Mal/Pyr) ratios were higher than in the control (Supplementary Figure S3).

Furthermore, metabolite responses to NO_3_^−1^ and Arg treatments differed in the case of citric acid and malic acid, glycine, Gly/Ser ratio, glycolic acid, ammonium and ornithine (Figure 8D, Supplementary Figure S3, Figure S4). Citric acid, malic acid and glycine levels increased significantly in NO_3_^−1^ treatments, whereas ammonium, ornithine and glycolic acid levels increased in Arg treatments (Figure 8D). Several amino acids also accumulated after the initiation of senescence, or before senescence in the highest NO_3_^−1^ and Arg treatments (Supplementary Figure S5) consistent with different roles of N forms in the process.

The Gly/Ser ratio fluctuated during autumn, but overall, it remained higher in NO_3_^−1^ treatments than in the control and was not significantly affected by Arg (Supplementary Figure S3). Metabolites affected by specific forms of N and the different responses of ornithine, glycine, Gly/Ser and C/N ratios indicated that inorganic NO_3_^−1^ and organic Arg fertilizations affected leaf metabolism at least partially through different pathways that could potentially be related to their different capacity to influence autumn senescence.

## Discussion

Throughout the world, N fertilization is used to increase plant yields in agriculture, and also sometimes in forestry. Excessive fertilization, however, can lead to nutrient leakage influencing water bodies, and the positive impacts of a yield increase from a CO_2_ perspective have to be balanced against the CO_2_ footprint of the production of commercial fertilizers (for a review, see e.g. Hedvall et al. 2014). One effect of N fertilization that has significant practical and economic consequences in boreal regions is that the overwintering capacity of trees, shrubs and some perennials may be compromised if fertilization is administered at the wrong time. Toca et al. (2018) reported that the effect of N availability on pine seedling frost tolerance differed between species and timing of the N treatment, and was not explained by timing of cessation of shoot elongation. Fertilization leading to growth stimulation late in the season influences the phenology of the tree and could hamper the development of hardiness, but the physiological understanding of this phenomenon is limited and hardly discussed in the scientific literature. Leaf senescence, in both trees and annuals, is connected to developmental age, but environmental conditions such as nutrient availability and drought affect the timing of senescence (Woo et al., 2013). We have developed *Populus* as a model system to study autumn senescence and have found that it is a precisely timed phenomenon which typically starts every year on almost the same date in a given genotype, grown at the same site (Keskitalo et al., 2005, Fracheboud et al., 2009). This precise timing is critical to allow for controlled remobilization of nutrients to avoid them getting lost due to sudden frost that can kill the leaf cells. Autumn senescence in trees in temperate and boreal regions should be seen from this perspective: a canopy that remains green longer in the autumn contributes to growth or storage but can jeopardize the retrieval of nutrients from the leaves. It seems that the timing of autumn senescence is genetically balanced as a trade-off between annual photosynthesis and nutrient balance, carbon *vs.* nitrogen. Boreal forest ecosystems are, in general, very N-poor (Högberg et al., 2017) and trees that do not senescence on time are at a disadvantage. Therefore, one could expect that trees would have developed the capacity to modulate autumn senescence in response to their nutrient, in particular N, status.

Addressing this issue poses experimental challenges. We made a few unsuccessful attempts to expose mature trees to various, controlled, fertilization doses in their natural environment to see if higher N levels could delay senescence, but faced several problems. We believed that one reason was that it was not possible to efficiently administer fertilizer to only one tree because in the rhizosphere there are functional connections between trees, perhaps even between different *Populus* clone individuals. Here, however, we have addressed this old question in a new way, not only by exposing trees to various nutrient regimes in the traditional way but also by “precision fertilization” through injecting trees in a clonal stand – sharing the same root system – with various amounts of different N species and monitoring the effect on leaf chlorophyll levels. To gain a better understanding of the underlying physiological processes, we also analyzed leaf metabolites after different fertilizations. We have to point out the complexity of these studies: in order to expose the trees to physiologically relevant conditions that induce autumn senescence in a very reproducible way, the studies were performed in the field. However, environmental conditions vary from year to year, but as we have performed these studies over several years, we believe that they reliably describe conditions to which the trees are adapted and acclimated.

We established that high levels of fertilization resulted in delay in the onset of senescence and reduced anthocyanin accumulation and that the delay could be achieved by high soil N availability without changing the availability of other nutrients. Furthermore, we used physiologically relevant N doses and “precision fertilization” to show that senescence onset could be delayed by NO_3_^−1^ whereas amino acids (Arg, Glu, Gln, or Leu) had no consistent effects. The delay in onset of senescence in the soil fertilization experiments was drastic – about a month – whereas the NO_3_^−1^ dose into the xylem caused less delay (ca. two weeks) but also this corresponds to approx. 10 % of the length of the growing period of these trees.

Major N-forms found in the xylem sap in *Populus* spp. are the amino acids, glutamine and asparagine, and nitrate, irrespective of N source in the growth media (e.g. Sauter and Cleve 1992; Siebrecht and Tischner 1999; Siebrecht et al. 2003; Millard et al. 2006). It has been suggested that during remobilization from the leaves, the main forms of transportable N are glutamine and asparagine, while N from the roots is transported in the form of nitrate (Siebrecht and Tischner 1999). The lowest concentrations of injected NO_3_^−1^ and amino acids were comparable to the physiological levels and the highest concentrations reached more than 30 times the levels typically found in the xylem sap of *Populus* spp. (Sauter and Cleve 1992; Millard et al. 2006; Hu and Guy, 2020).

The fact that the addition of amino acids (Arg, Glu, Gln, and Leu) in the doses administered here did not affect autumn phenology has several implications. First and foremost, as nitrogen in forest soils is mainly present in organic forms (Schulten et al., 1998, Yu et al., 2002, Andersson et al., 2004, Inselsbacher and Näsholm 2012), the phenology of the tree will be better “buffered” against changes in soil nitrogen content, which could be important as the N content could change a lot during the lifespan of a tree while the key factor for local adaptation of phenology – the climate – is likely to change less. Secondly, as delayed autumn phenology could jeopardize the survival of the tree over the winter, late-season fertilization with Arg could provide an attractive alternative to traditional inorganic fertilizers either in tree nurseries or in forest plantations.

We also aimed to identify metabolic processes associated with *Populus* leaf senescence in relation to N status and N forms. Soil N fertilization increased pre-senescence chlorophyll levels considerably, whereas “milder” fertilization directly into the xylem stream affected neither pre-senescence chlorophyll levels nor N levels in the leaves, indicating that the amount injected was within the physiological levels of N and, in the case of mature trees, the injected N form was more important than the absolute N level for modulating senescence onset in autumn.

In general, soil N fertilization as NH_4_NO_3_ or injection as KNO_3_ in the xylem stream affected leaf metabolite profiles more than did Arg injection. Metabolite responses to soil and xylem N fertilization with NH_4_NO_3_ or KNO_3_, respectively, were, however, different. This may have been due to many factors, including the different N forms and the application methods, since the uptake of different forms of N is regulated by the root system. In addition, they can be linked to the different levels used in relation to physiological levels, as well as to the capacity of soil and xylem fertilization to affect the internal N status of trees (strong effect in chlorophyll levels after soil fertilization and little or no effect in chlorophyll and nitrogen levels after xylem fertilization).

The observed shifts in the leaf metabolite profiles and the peaking metabolite levels during the experiment indicated that the N compounds injected into the xylem stream reached the leaves and influenced leaf metabolism irrespective of the form. However, it is possible that a significant fraction of the added N ended up elsewhere, such as in buds, twigs, and the bark, in storage proteins (bark storage protein, BSP, is a major storage constituent of *Populus* bark) acting as N sinks (Sauter et al., 1989). Nevertheless, we believe that the injection system for delivery of known amounts of solutes or labelled metabolites into *Populus* trunks could potentially be used to answer fundamental questions about N transport and storage during autumn. Our results demonstrate that N availability affected many cellular functions in the leaves, including adjustment of metabolic pathways involved in C and N metabolism, respiration and photorespiration. Notably, senescence under low N availability (LN) did not coincide with the accumulation of soluble sugars, which has been widely reported in other plant species (e g Balazadeh et al., 2014; Have et al., 2017). This may be a consequence of the presence of strong sinks for carbohydrates in *Populus* during autumn, for example carbon-based secondary metabolites such as anthocyanins or storage carbohydrates such as starch, alleviating soluble carbohydrate accumulation in the leaves. It has been reported that carbohydrates with a presumed signaling role, for example trehalose-6-phosphate (T6P), accumulate under N deficiency in barley, coinciding with a minor accumulation of other soluble carbohydrates (Fataftah et al., 2018). The levels of glucose, fructose, and galactose were higher in the leaves of HN saplings than of LN saplings (Figure 7). These results suggest that carbohydrate metabolism in response to N availability and during autumn senescence in *Populus* is different to the often-studied herbaceous model species.

During the senescence process, the respiration rate increases as the TCA cycle and oxidative phosphorylation in mitochondria become the main energy-producing pathways to fuel the remobilization of nutrients (Chrobok et al. 2016; Law et al. 2018). Both soil and xylem fertilization treatments affected the levels of TCA cycle intermediates, but in different ways. An increase in free amino acid levels and the Gln/Glu ratio (indicator of ammonium assimilation) is a typical response to high N availability (Urbanczyk-Wochniak and Fernie 2005; Schlüter et al. 2012) and was also observed here under high soil N availability (Figure 6). Often the levels of TCA cycle intermediates also increase (Urbanczyk-Wochniak & Fernie 2004; Schlüter et al. 2012), and the results in the xylem fertilization treatments with NO_3_^−1^ and Arg were in line with this (Figure 8). Contrary to our expectations, although the amino acid levels were higher in the soil fertilization experiment, the levels of TCA cycle intermediates were lower in the HN than the LN regime. Maybe under certain conditions the levels of TCA intermediates and derived amino acids can be uncoupled, presumably as the organic acids are shuttled between cellular compartments to transfer reducing equivalents. The balance between amino acid biosynthesis and degradation under different N regimes may also influence the TCA cycle.

The partially different responses of leaf metabolite levels and the C/N ratio in NO_3_^−1^ and Arg treatments also indicated that they induced different adjustments of cellular metabolism. These different responses can be related to the nitrogen forms used in the experiments, since the assimilation of nitrate and ammonium (released by Arg catabolism) has different metabolic requirements for reducing power and carbon skeletons. Accordingly, the Arg treatment increased ammonium levels as well as ornithine levels in the leaves, indicating that arginine was efficiently catabolized. NO_3_^−1^ and NH_4_^+^ are known to modify major metabolic pathways such as glycolysis (Masakapalli et al., 2013), amino acid biosynthesis, TCA cycle (Vega-Mas et al. 2019) and photorespiration (Guo et al. 2007) differently. The main differences between NO_3_^−1^ and Arg treatments were seen in the levels of glycine and the Gly/Ser ratio (a photorespiration indicator) that were higher in the NO_3_^−1^ treatments than in the control, and not affected by Arg (supplementary Figure S3). This is in line with the reports showing that NO_3_^−1^ induces photorespiration because its assimilation requires reducing equivalents (Guo et al., 2005; Guo et al. 2007). However, soil N fertilization with ammonium nitrate had the opposite effect on the Gly/Ser ratio as it was lower in the HN than in the LN treatment, further indicating that the responses could depend on the regulated uptake of N form by the root system and/or the relative levels of NO_3_^−1^ and NH_4_^+^. The TCA cycle is regarded as being the critical pathway that connects N assimilation, respiration and photorespiration (Foyer et al. 2011). Our study demonstrated that all of these pathways were affected by N treatments, but the metabolic adjustments and the capacity to delay senescence depended on the concentration used as well as the form of N applied. How the tightly interlinked amino acid metabolism, TCA cycle and photorespiration pathways interact during autumn senescence in response to the ways of NO_3_^−1^ application and levels is an intriguing topic for further study.

## Conclusions

In conclusion, we have shown that nitrogen fertilization has the capacity to delay autumn leaf senescence, and the effect is more pronounced for inorganic nitrate than for organic amino acids. Since amino acid fertilization did not affect autumn phenology of trees even at quite high concentrations, arginine fertilizer could be a potential alternative to typical inorganic fertilizers in tree nurseries and forest plantations late in the season. However, the optimal levels of organic N fertilizer that do not cause any adverse effects on tree physiology, such as on cold acclimation or frost tolerance, remain to be determined in the future. The effects of the different fertilization regimes turned out to be complex, but soluble carbohydrate accumulation was not associated with autumn senescence, even under our low N regime. Instead, the tightly interlinked metabolic pathways of amino acid metabolism, the TCA cycle and photorespiration appeared differently affected by arginine and NO_3_^−1^, in soil treatments and when injected, suggesting that the direct molecular signaling of NO_3_^−1^ and redox metabolism could account for the regulation of autumn senescence in *Populus*. Overall, the form of applied N fertilization appeared to be more important than the N status of the tree for influencing autumn senescence.

## Material and Methods

### Plant material and experimental setups

#### Fertilization experiment in a common garden

Fertilization was performed on an experimental plot adjacent to the SLU greenhouse on Umeå University campus in autumn 2008. Thirteen *Populus* from four different genotypes from the Swedish *Populus* (SwAsp) collection (Luquez et al. 2007) were used for the experiment: genotype 88 from the Dorotea population and genotypes 91, 99 and 100 from the Umeå population, and they were therefore locally adapted. The trees were planted in sandy soil with limited nutrient levels, the pots were left around the root systems although roots were allowed to grow out through the holes in the pots into the surrounding soil. All plants received equal doses of garden fertilizer “Blåkorn” containing macro and micronutrients (N [NH4NO3]-P-K 12:3:13, Bayer AB) from the beginning of summer until July 7th 2008, when fertilization was stopped on half the plot (Low fertilizer; LF) and continued on the other half (high fertilizer; HF). The HF treatment received approx. 300 kg of N/ha. The two parts of the plot were separated by two rows of trees, not included in the experiment, to ensure that the unfertilized trees did not access the fertilizer.

#### Soil fertilization experiment with inorganic nitrogen

The hybrid *Populus* plants (genotype T89, *Populus tremula* × *tremuloides*) were obtained from tissue culture and were potted in 3 liters of mull soil containing low nutrient levels and 1.1 g/L of total N (mainly in the form of organic matter). Plants were fertilized with 100 ml of modified nutrient solution including N and covered with a plastic bag for two weeks, after which they were fertilized with 50 ml of modified nutrient solution (pH 5.8) twice a week until late autumn. The nutrient solution in the low nitrogen treatment (LN) contained 3 mM K_2_SO_4_, 3 mM CaSO_4_, 3 mM KH_2_PO_4_, 2 mM MgSO_4_, 100 μM Fe-EDTA, 100 μM K_2_HPO_4_, 50 μM MnSO_4_, 50 μM H_3_BO_3_, 20 μM Na_2_MoO_4_, 10 μM ZnSO_4_, 3 μM CuSO_4_, and 0.2 μM CoSO_4_ and in the high nitrogen treatment (HN) the solution contained an additional 9 mM of NH_4_NO_3_. The plants were kept in a greenhouse with a 18/6 h photoperiod under 200 μmol m^−2^ s^−1^ light and 20/15°C (day/night). After 40 days, the plants were transferred to natural conditions outside the greenhouse (on August 22^nd^ 2018). The chlorophyll content of four leaves from the lower or the upper parts of the shoot was measured (*n*=6 plants). The lower leaves were sampled from 4-6 trees for metabolite analysis on four dates always at noon, frozen in liquid nitrogen and stored at −80 °C.

### Xylem fertilization experiments with inorganic and organic nitrogen

Much *Populus* propagation in the natural environment is through clonal replication, where several suckers emerge from the same root, resulting in mature trees sharing the same root system. The field fertilization plots were chosen from large stands of *Populus* (10 trees or more). Four stands were selected, and 10 trunks were randomly selected within each stand (one per treatment and two for controls). However, one of the stands – growing next to an agricultural field – turned out to have very high chlorophyll levels, much higher than we ever recorded for any aspen tree grown in the field. As this was probably a consequence of the stand being influenced by fertilization of the adjacent field, it was excluded from further analysis. The procedure was a modification of the passive liquid injection method described by authors including Swanston and Myrold (1998) and Proe et al. (2000). A hole (diameter 15 mm, depth ca. 5 cm) was drilled through the bark into the xylem. The hole was filled with a rubber cork and sealed with silicon glue that covered the phloem but not the xylem. The hole was subsequently filled with water by inserting a syringe on the top of the cork to let the air out as the water was injected. A refillable infusion bag was suspended on the tree and connected to the hole by a syringe. The bags were filled with 2 L of the solution and replaced every week. On eight occasions, from July 26^th^ to September 13^th^, 2011, different amounts of nitrate in the form of KNO_3_ or Arg (arGrow® Support (N-70g/L)) in solution were added to the bags right before they were attached to the trees. We gave four different doses of N on eight occasions over the season, in total 4, 8, 20, or 60 g of N per trunk were injected (hereafter referred to as N4, N8, N20, and N60 for NO_3_^−1^, and A4, A8, A20 and A60 for Arg treatments). Five leaves were sampled per trunk twice a week from mid-July until mid-October. The samples were pooled for metabolite, carbon and nitrogen analyses and, at the same time, chlorophyll content was measured. Samples were frozen in liquid nitrogen and stored at −80 °C.

In 2018, in the second injection experiment, trees were injected on nine occasions from August 14^th^ to September 25^th^. Solutions were prepared and added to the bags right before application on the trees, including inorganic (KNO_3_; 5 g of total injected N over the season; pH 6) or organic N (Glu, Gln, or Leu; 2.5, 5, and 2.5 g of total injected N over the season, respectively; pH 6). The trunks were injected with 1L of the solution every week in August, and twice a week in September. Chlorophyll content of leaves was measured for 10 leaves from each trunk through the experiment. In both xylem fertilization experiments, control stems were injected with a similar volume of distilled water.

### Pigment measurements and the onset and rate of senescence

In all experiments, the chlorophyll content index (CCI) was measured with a CCM200+ chlorophyll content meter and the anthocyanin content index (ACI) was measured with an ACM-200+ anthocyanin content meter (Opti-Sciences, Hudson, USA). To determine the dates of onset and termination of senescence, and senescence rate, the CCI values were fitted with a non-linear function with OriginPro (version 9, 2019, OriginLab Corporation, Northampton, MA, USA) as described in Lihavainen et al. (2020).

### Carbon and nitrogen content

In the xylem fertilization experiment in autumn 2011, involving nitrate (KNO3) and arginine, carbon (C) and nitrogen (N) contents of leaves were determined from samples collected on 12 dates during autumn. Leaves were homogenized in liquid nitrogen with mortar and pestle, and leaf powder (10 mg dry weight) was analyzed with a Thermo Fisher Scientific flash EA 1112 NC Soil Analyser. The data were used to calculate the carbon-to-nitrogen ratio (C/N), and the N content (mg/cm^2^) in the leaves before and after senescence was used to calculate the relative amount of remobilized N. The effects of time, fertilization treatment (nitrate or arginine) and their interaction with time on the C/N ratio were tested with two-way ANOVA, and the treatment effects on nitrogen levels and remobilization with one-way ANOVA, a *P*-value <0.05 was considered significant (IBM SPSS Statistics, version 25).

### Metabolite analyses

In the soil fertilization experiment with low (LN) or high nitrogen (NH4NO3) treatment (HN), metabolite analyses were performed at four time points during autumn, i.e. on August 16^th^, 22^nd^ and 29^th^, and on September 5^th^ 2018. Leaves were pooled from six trees for each treatment/time, homogenized in liquid nitrogen with mortar and pestle, then 10 mg of leaf powder were used in the analysis. Extraction and derivatization were performed as described in Gullberg et al. (2004). The GC-MS system consisted of a GC PAL autosampler (CTC Analytics, Zwingen, Switzerland) combined with a column oven (7890A Agilent Technologies, USA) and Pegasus HT GC-MS/QTOF (Leco, St. Joseph, MI, USA) with electron ionization of 70 eV. Each sample (1 μl) was injected in 20:1 split mode into a metal liner with deactivated wool (5.2 × 6.3 × 78.5 mm, Restek). The inlet temperature was set to 260°C and helium flow in the column (Agilent J&W DB-5MS Ultra Inert 30 m, ∅ 0.25 mm, 0.25 μm) was kept constant at 1 ml min^-1^. The temperature of the column was maintained at 70°C for 2 min, and then increased by 30°C min^-1^ to 200°C, by 5°C min^-1^ to 220°C, and by 15°C min^-1^ to 320°C, where the temperature was kept for 4 min. Transfer line and source temperatures were 270°C and 200°C, respectively. Scans were recorded at a rate of 20 spectra s^-1^ in a mass range of 50-800 m/z. The contents of soluble carbohydrates were quantified (mg/g FW) based on the standard curves of corresponding reference compounds.

In the xylem fertilization experiment, untargeted and targeted metabolite analyses were performed for samples collected on 12 dates in autumn 2011 for the leaves of control trees, and in the leaves of trees injected with different amounts of nitrate (KNO_3_) and arginine. The samples (approx. 10 mg FW) were extracted, derivatized and analyzed with gas chromatography-mass spectrometry (GC-MS) as described in Gullberg et al. (2004). Targeted amino acid analysis was performed using a Waters UPLC Amino Acid Analysis Solution with ACQUITY UPLC® (Ultra performance liquid chromatography) System combined with a tunable UV (TUV) detector (Waters, Milford, MA, USA). Samples were derivatized with an AccQ•Tag™ Ultra Derivatization kit and the analytes were separated with an AccQ•Tag™ Ultra Column (ACQUITY UPLC BEH C18, 2.1 × 100 mm, 1.7μm) at a flow rate of 0.6 ml/min. The amino acid contents were quantified (nmol/g FW) based on the standard curves of corresponding reference compounds.

In both experiments, in-house software developed by the Swedish Metabolomics Centre (RDA software version 2016.04) was used to process the GC-MS data (netCDF files). Metabolite peak areas were normalized with the peak area of internal standards and the samples’ fresh weight (FW). All statistics were performed with normalized data. Metabolites (86 metabolites in both data sets) were annotated based on retention index (RI based on alkane series) and mass spectra that were compared to an in-house library (reference standards) and databases (NIST, Golm metabolome database GMD, Fiehn library).

### Data processing and statistics

Statistical analyses were performed with log10-transformed data with unit variance scaling. Metabolite data were analyzed with principal component analysis (PCA, Simca P+ version 15, Umetrics, Sweden) to detect general variation in the data sets in both soil and xylem fertilization experiments. When testing the effects of time or treatments on individual metabolite levels, *P*-values were adjusted with a false discovery rate (FDR) for multiple comparisons. In all statistical analyses, a *P*-value <0.05 was considered significant.

In the soil fertilization experiment, the N treatment effect on individual metabolite levels and metabolite ratios was tested with a t-test (MetaboAnalyst, Chong et al. 2018). A volcano plot was produced to display the significance versus the magnitude of change (log_2_ fold change cut off 1.0) in the metabolite levels between the two nitrogen treatments.

In the xylem fertilization experiment, the initial chlorophyll levels and metabolite profiles differed between the tree stems before fertilization started on July 26^th^ in 2011. Therefore, the metabolite levels were normalized based on the pre-fertilization levels on July 4^th^ to reveal changes in response to treatments during autumn. The effects of time and fertilization treatment on metabolite levels were tested with two-way ANOVA, the effects of nitrate and arginine treatments were tested separately (MetaboAnalyst). The time×treatment interaction term was not significant in either of the xylem fertilization forms, and thus it was left out of the models. The interaction term was kept in the model when testing the time and treatment effects on the metabolite ratios. Since a clear separation of xylem fertilization treatments from the control was observed in the PCA, orthogonal projection to latent structures discriminant analyses (OPLS-DA, Simca P+) were performed to identify metabolites that accounted for most for the separation of treatments. OPLS-DA models were produced separately to compare metabolite profiles between control and nitrate treatments and between control and arginine treatments. Both of the models were of good quality (Control vs. Nitrate: R^2^ 0.856, Q^2^ 0.799; Control vs. Arginine: R^2^ 0.959, Q^2^ 0.862) and significant (CV-ANOVA *P*-value <0.001). The metabolites with p(corr)-values of >|0.3| and VIP-scores (Variable Importance to Projection) >1.0 were regarded as the most important ones to account for the separation of the treatments from the control.

## Supplementary Data

Supplementary Figure S1. Chlorophyll content in trunks in three aspen stands that were injected with water (Control, C), nitrate (KNO_3_, N4-N60) or arginine (A4-A60) in autumn 2011.

Supplementary Figure S2. Chlorophyll content index in pre-senescence leaves in xylem fertilization experiment in autumn 2011.

Supplementary Figure S3. Metabolite ratios as affected by xylem fertilization treatments during autumn 2011 in aspen leaves.

Supplementary Figure S4. Metabolite levels as affected by xylem fertilization treatments during autumn 2011 in aspen leaves.

Supplementary Figure S5. Heatmap of leaf amino acid levels in aspen in xylem fertilization experiment.

### Supplementary tables

Supplementary Table S1. The effect of soil fertilization (HN/LN) on metabolite levels in aspen leaves during autumn 2018.

Supplementary Table S2. The effects of time and xylem fertilization treatment with nitrate (KNO3) on individual metabolite levels, metabolite ratios and carbon-to-nitrogen (C/N) ratio in aspen leaves during autumn 2011.

Supplementary Table S3. The effects of time and xylem fertilization treatment with arginine on individual metabolite levels, metabolite ratios and carbon-to-nitrogen (C/N) ratio in aspen leaves during autumn 2011.

## Acknowledgments

We thank the Swedish Metabolomic Centre (SMC) for their help in metabolite analyses and data processing.

